# The effect of drought and intercropping on chicory nutrient uptake from below 2 m studied in a multiple tracer setup

**DOI:** 10.1101/562876

**Authors:** Camilla Ruø Rasmussen, Kristian Thorup-Kristensen, Dorte Bodin Dresbøll

## Abstract

**Aims:** We tested if chicory acquires nutrients from soil layers down to 3.5 m depth and whether the deep nutrient uptake increases as a result of topsoil drought or topsoil resource competition. We also tested whether application of the trace elements Cs, Li, Rb, Sr, and Se, as tracers result in similar uptake rates.

**Methods:** The methodological tests were primarily carried out in a pilot experiment where the five tracers were applied to 1 m depth in lucerne and red beet grown in tube rhizotrons. The dynamics of deep nutrient uptake in chicory was studied in large 4 m deep rhizoboxes. A drought was imposed when roots had reached around 2 m depth.

**Results:** Chicory acquired tracers applied to 3.5 m depth, but we found no compensatory tracer uptake with depth during drought. We found some indications of a compensatory tracer uptake from 2.3 and 2.9 m depth in intercropped chicory. Application of equimolar amounts of trace elements resulted in similar excess tracer concentrations within species.

**Conclusion:** Chicory acquires nutrients from below 3 m but does not increase deep nutrient uptake as a response to limited topsoil nutrient availability.

## Introduction

The key advantage of deep roots growing below 1 m depth or even further down is classically perceived to be the access to larger water pools, contributing to water uptake both under dry and well-watered conditions (Gregory et al. 1978; Nepstad et al. 1994; Jackson et al. 1999; Elliott et al. 2006; Maeght et al. 2013). Deep roots have also shown to play a significant role in the uptake of nitrogen (N) deep in the soil profile (Buresh and Tian 1997; Thorup-Kristensen 2001). N is mobile in the soil and surplus precipitation is causing N to leach (van Noordwijk M 1989; Jobbágy and Jackson 2001) making downward root expansion more valuable than increasing root densities in an already occupied soil volume (McMurtrie et al. 2012).

Efficient utilisation of less mobile nutrients requires a higher root density, and uptake is therefore generally associated with topsoil root activity. The higher organic matter content and fertilisation in cropping systems, ensure the release of nutrients in the topsoil. However, uptake of nutrients deeper in the soil profile might be an overlooked resource at some places. Considerable amounts of plant available nutrients can be present below 1 m depth (Stone and Comerford 1994; Jobbágy and Jackson 2001; McCulley et al. 2004). Lehmann (2003) stressed that despite the usually lower relative root activity in the subsoil compared to the topsoil, the large volume of subsoil in comparison to the mostly narrow band of topsoil represents an important resource for nutrient uptake.

Even though the highest concentration of nutrients is often found in the topsoil, the availability of these nutrients decreases when the topsoil dries out. This occurs especially in semi-arid environments (reviewed by da Silva et al. 2011a), and nutrient uptake from deeper soil layers becomes increasingly important here (Lehmann 2003; Ma et al. 2009). Moving towards more sustainable cropping systems with lower nutrient inputs might also increase the importance of utilising available nutrients deep in the profile (Lynch 2007). In addition, it is not unreasonable to assume that increasing the average rooting depth in cropping systems for various purposes such as drought tolerance, carbon sequestration, soil fertility, and reduced soil compaction will itself increase nutrient deposition deeper in the profile, as it is seen for carbon deposition (Rasse et al. 2005; Kell 2011).

Methods using trace elements as tracers have shown to be valuable in detecting temporal and spatial patterns of nutrient uptake in plants and plant mixtures (Sayre and Morris 1940; Martin et al. 1982; Casper et al. 2003; Hoekstra et al. 2014). Substances qualify as tracers if they naturally occur in very low quantities, are chemically equivalent to the nutrients of interest and are non-toxic for plants in concentrations that are readily determinable (Pinkerton and Simpson 1979). Strontium (Sr) is physiologically an analogue to calcium (Ca) and is absorbed following the plant’s metabolic requirements for Ca (Kabata-Pendias 2011). Lithium (Li), cesium (Cs) and rubidium (Rb) all form monovalent cations that appears to share the K+ transport carrier (Isaure et al. 2006; Valdez-Aguilar and Reed 2008; Kabata-Pendias 2011). Selenium (Se) is mainly taken up as 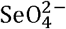, and is thus an analogue to sulfur (Terry et al. 2000)

Despite the different uptake transport carriers, the cation trace elements are also used interchangeable simply as root activity tracers, potentially allowing injection into multiple depths of the same plot (Fitter 1986; Carlen et al. 2002). Compared to using one tracer in each plot this reduces the between-plot variability, but few studies have rigorously tested the feasibility of the strategy (Hoekstra et al. 2014).

Most studies making use of trace elements as tracers are focusing on the upper 1 m of soil or even shallower. To the author’s knowledge only three studies are placing trace elements deeper, of which two are focusing on eucalyptus trees (da Silva et al. 2011a; Bordron et al. 2019) and only one on an agricultural crop, that is lucerne (Fox and Lipps 1964). All three studies documented significant uptake of Sr, and the eucalyptus studies also of Rb at 3 m depth, demonstrating the feasibility of the method.

In the field, placing tracer at great depths can be challenging compared to shallower placement, limiting the number of sites of placement. Also, the risk of damaging the root system increases with the depth of placement as fewer roots are usually reaching the greater depths. Thus causing damage to these critical structures for deep nutrient acquisition can alter the plant behaviour. However, decreasing the number of placement sites causes an increase in variation in uptake (Fox and Lipps 1964; Hoekstra et al. 2014), which presumably increases with depth as the root system is less dense. It might in some experiments be relevant to be able to remove the tracer-infused soil again, allowing repeated placement of tracers at the same site at different times. If this is the case, the tracer cannot be distributed evenly, and the number must be limited.

In this study, we used Cs, Li, Rb, Se and Sr to test the possibility of using the trace elements interchangeable as tracers and to test the feasibility of placing the tracers via ingrowth cores, letting roots grow into tracer infused soil placed at various depths. We also examined the effect of topsoil drought and intercropping on deep nutrient uptake in chicory. More specifically, we tested the hypotheses that 1) application of the less soil mobile trace elements Cs, Li, Rb and Sr as tracers result in similar uptake rates, whereas the mobile trace element Se is taken up at a higher rate, 2) placing trace element tracers in ingrowth cores is a feasible method to avoid contamination of the soil, enabling repeated use of tracers in experimental setups, 3) chicory acquire both mobile and less mobile nutrients from soil layers down to 3 m, and 4) the deep nutrient uptake increases as a result of topsoil drought or topsoil resource competition.

To test hypothesis 1) we set up a pilot study where we applied the five trace element tracers to 1 m depth in lucerne and red beet grown in 1 m tall tube rhizotrons. To test hypothesis 2) and 3) we grew chicory (Cichorium intybus L.) as a sole crop and intercropped with the two shallow-rooted species ryegrass (Lolium perenne L.) and black medic (Medicago lupulina L.) in 4 m tall rhizoboxes. We allowed the roots to reach around 2 m depth before imposing a drought, as our focus was on the potential of deep roots to acquire nutrients and not on deep root growth during drought. Details on soil water content during the experiment and the implications of the depth of plant water uptake has been reported in Rasmussen et al. (2019a, preprint) together with repeated observations of root development obtained from the transparent surfaces of the rhizoboxes.

## Methods

Both experiments were placed outside and situated at University of Copenhagen, Taastrup, Denmark (55°40’8.5”N 12°18’9.4”E).

### Pilot experiment

We grew lucerne *(Medicago sativa L.)* and red beet *(Beta vulgaris L.)* in 1 m tall transparent acrylic glass tubes with a diameter of 150 mm in 2 x 4 replicates for pre- and post-tracer harvest. The tubes were placed on wooden frames outside and wrapped in white non-transparent plastic to avoid light exposure of soil and roots.

Growth medium was a sandy loam soil from a research farm in Taastrup, Denmark (Table 1), belonging to University of Copenhagen. The bottom 0.75 m of the tubes was filled with soil from the lower horizon (0.2–0.6 m) and the top 0.25 m of the tubes was filled with soil from the upper horizon (0–0.2 m). We sieved the soil through a 15 mm sieve and mixed it thoroughly before filling it into the tubes compacting it for each 0.03 m to a bulk density of approximately 1.33 g cm^-3^. A plastic net covered the bottom of the tubes, and they were placed on a wick to allow water to drain. Seeds of lucerne and red beet were sown in 40 × 40 mm pots in a greenhouse and transplanted into the tubes on 20 June 2015, one week after sowing. After transplanting, we fertilized the plants with NPK 5-1-4 fertilizer equivalent to 50 kg N ha^-1^. During the experiment, plants were exposed to precipitation and were further irrigated using ceramic cups that ensured slow water infiltration when necessary.

On 6 August, we placed a 2 l plastic pot filled with subsoil under each tube of the post-tracer harvest replicates. Roots could grow through the plastic net and into the pots. In each pot, we had applied Cs, Li, Se, Rb, Sr to trace root activity. We did this by dissolving CsNO_3_, LiNO_3_, Na_2_SeO_4_, RbNO_3_ and Sr(NO_3_)_2_ in water to a concentration resulting in an application of 3 g, 0.2 mg, 2 g, 2 g, 2 g rrf^2^, equivalent to 23, 29, 0.025, 23 and 23 mmol m^2^ respectively, and mixing it into the soil. We harvested aboveground biomass of the pre-tracer replicates 10 August and the post-tracer harvest replicates 1 September. All biomass samples were dried at 80°C for 48 hours. Samples were ground and the concentration of the trace elements was determined on an ICP-MS (Thermo-Fisher Scientific iCAP-Q; Thermo Fisher Scientific, Bremen, Germany) at University of Nottingham, School of Biosciences laboratory. We calculated excess tracer concentration in aboveground biomass as the increase in concentration of each of the trace elements from the pre-tracer harvest to the post-tracer harvest.

Root development was documented every week by photographing the roots growing at the soil-tube interface, with an Olympus Tough TG-860 compact camera. We took 10 photos to cover the tube from top to bottom and at four different positions at each height. In total, the images covered 40% of the tube surface. We recorded the roots using the line intersects method (Newman 1966) modified to grid lines (Marsh 1971; Tennant 1975) to calculate root intensity, which is the number of root intersections m-1 grid line in each panel (Thorup-Kristensen 2001). The grid we used was 10 × 10 mm.

### Experimental facility – main experiment

We conducted the main experiment in a semi-field facility and repeated it for two consecutive seasons, 2016 and 2017. We grew the crops in 4 m deep rhizoboxes placed outside. The boxes were 1.2 x 0.6 m and divided lengthwise into an east- and a west-facing chamber. On the wide east- and west-facing sides of the boxes, 20 removable panels allowed us to access to the soil column at all depths. All sides of the boxes were covered in white plates of foamed PVC of 10 mm thickness.

We used field soil as growth medium. We filled the bottom 3.75 m of the rhizoboxes with subsoil taken from below the plow layer at Store Havelse, Denmark (55°89’83.9”N, 12°06’52.8”E, Table 1), and the upper 0.25 m with a topsoil mix of clay loam and sandy loam half of each, both from the University’s experimental farm in Taastrup, Denmark. Soil bulk density was 1.6 g m^3^, which is close to field conditions for this soil type. We filled the boxes in August 2015 and did not replace the soil during the two experimental years. Rainout shelters constructed of tarpaulin were mounted to control precipitation. Drip irrigation with a controlled water flow of 15 mm hour^-1^ was installed in each chamber. For a thorough description of the facility, see Rasmussen et al. (2019a, preprint).

**Table 1.** Main characteristics of the soil used in the tubes and the rhizoboxes

| Experiment | Depth (m) | Organic matter*<sup>1</sup><br>(%) | Clay (%)<br><0.002 mm | Silt (%)<br>0.002-0.02 mm | Fine sand (%)<br>0.02-0.2 mm | Coarse sand (%)<br>0.2-2.0 mm | pH**<sup>2</sup> |
| --- | --- | --- | --- | --- | --- | --- | --- |
| Tubes | 0.00-0.25 | 2.3 | 7 | 6 | 55 | 30 | 6.8 |
|  | 0.25-0.75 | 0.6 | 6 | 5 | 57 | 32 | 6.4 |
| Rhizoboxes | 0.00-0.25 | 2.0 | 8.7 | 8.6 | 46.0 | 35.0 | 6.8 |
|  | 0.25-4.00 | 0.2 | 10.3 | 9.0 | 47.7 | 33.0 | 7.5 |

\*<sup>1</sup> Assuming that organic matter contains 58.7 % Carbon.
\*\*<sup>2</sup> pH = Reaction Number (Rt) – 0.5. Measured in a 0.01 M CaCl<sub>2</sub> suspension, soil:suspension ratio 1:2.5.

### Experimental design

We had two treatments in 2016 and four treatments in 2017. In both years we grew chicory *(Cichorium intybus L*, 2016: Spadona from Hunsballe frø. 2017: Chicoree Zoom F1 from Kiepenkerl) in monoculture under well-watered (WW) and drought stress (DS) conditions. We chose to work with a hybrid salad cultivar the second year to reduce the variation among plants in size and in development speed seen in the forage type used the first year. In 2017, we also grew chicory intercropped with either ryegrass *(Lolium perenne L.)* or black medic *(Medicago lupulina)*, both in a WW treatment. In 2016, we transplanted four chicory plants into each chamber and in 2017, we increased the number to six in order to reduce within-chamber variation. For the two intercropping treatments in 2017, we transplanted five plants of ryegrass or black medic in between the chicory plants.

For the 2016 season, chicory plants were sown in May 2015 in small pots in the greenhouse and they were transplanted into the chambers 30 September. Despite our attempt to compact the soil, precipitation made the soil settle around 10 % during the first winter. Therefore, 29 February 2016, we carefully dug up the chicory plants, removed the topsoil, filled in more subsoil to reach 3.75 m, before filling topsoil back in, and replanting the chicory plants. A few chicory plants did not survive the replanting and in March, we replaced them with spare plants sown at the same time as the original ones and grown in pots next to the rhizoboxes. In 2017, we sowed chicory in pots in the greenhouse 11 April and transplanted them to the chambers 3 May (Table 2). Chicory is perennial, it produces a rosette of leaves the first year and the second year it grows stems and flowers. Due to the different sowing strategies, we worked with second-year plants in 2016 and first-year plants in 2017.

**Table 2.** Timeline of the experiments in 2016 and 2017

|  | 2016 | 2017 |
| --- | --- | --- |
| Sowing | May 2015 | 11 April |
| Transplanting | 29 February | 3 May |
| Onset of drying out | 26 June | 13 July |
| Insertion of ingrowth cores | 12 July | 15 August |
| Removal of ingrowth cores | 28 July | 11 September |
| Cut of aboveground biomass | 28 July | 11 September |
| Onset of drying out | 12 August |  |
| Insertion of ingrowth cores | 23 August |  |
| Removal of ingrowth cores | 22 September |  |
| Cut of aboveground biomass | 22 September |  |

We grew all treatments in three randomized replicates. The soil inside the six chambers not used for the experiment 2016 but included in 2017 had also sunken during the 2015/2016 winter and we used the same procedure of filling in more soil for these chambers.

In 2016, we had two experimental rounds using the same plants. For the 1^st^ round, we started drying out the DS treatments 26 June and ended the round by cutting aboveground biomass 28 July. Hereafter we let the plants regrow until we again started drying out the DS treatments 12 August for the 2^nd^ round. We swopped treatments so that chambers used for DS treatments in the 1^st^ round where used for WW treatments in the 2^nd^ round and vice versa. We pruned the plants at 0.5 m several times between 24 May and 12 July 2016 to postpone flowering and induce leaf and root growth. In 2017, we only had one round, starting the drying out on 13 July 2017. We fertilized all chambers with NPK 5-1-4 fertilizer equivalent to 50 kg N ha^-1^ on 1 April and 21 June in 2016 and again on 29 July before the 2^nd^ round in 2016. In 2017, we fertilized all chambers 3 May and 1 June following the same procedure.

In the DS treatment, we stopped irrigation and mounted the rainout shelters. In WW treatments including the intercropping treatment in 2017, we irrigated regularly. In 2017, we chose to supply the same amount of water in all chambers, apart from in the DS treatment, which led to different levels of soil water content due to differences in evapotranspiration. In 2017, two chambers were accidentally over-irrigated mid-June 2017 and we re-fertilized them 16 June.

### Soil water content

We installed time-domain reflectometry sensors (TDR-315/TDR-315L, Acclima Inc., Meridian, Idaho) at two depths to measure volumetric water content (VWC) in the soil. In 2016, the sensors were installed at 0.5 and 1.7 m depth. In 2017, the sensors were installed at 0.5 and 2.3 m depth. Soil water content was recorded every 5 min in 2016 and every 10 min in 2017 on a datalogger (CR6, Campbell Scientific Inc, Logan, Utah).

### Ingrowth cores and tracer application

We made ingrowth cores out of PVC pipes with a diameter of 70 mm and a length of 250 mm. We drilled four holes, 50 mm in diameter on each side of the cores. When placed horizontally with holes facing up- and downwards, the holes allowed roots to grow into the cores from the upper side and out below the cores (Figure 1). In 2016, we filled the cores with 1900 g of moist soil. We later dried three samples from each soil batch and calculated that in all cases soil bulk density was between 1.55 and 1.59 m^3^ and volumetric water content was between 27.4 and 29.9 %. In 2017, we followed the same procedure but did not check the water content.

**Fig. 1.**
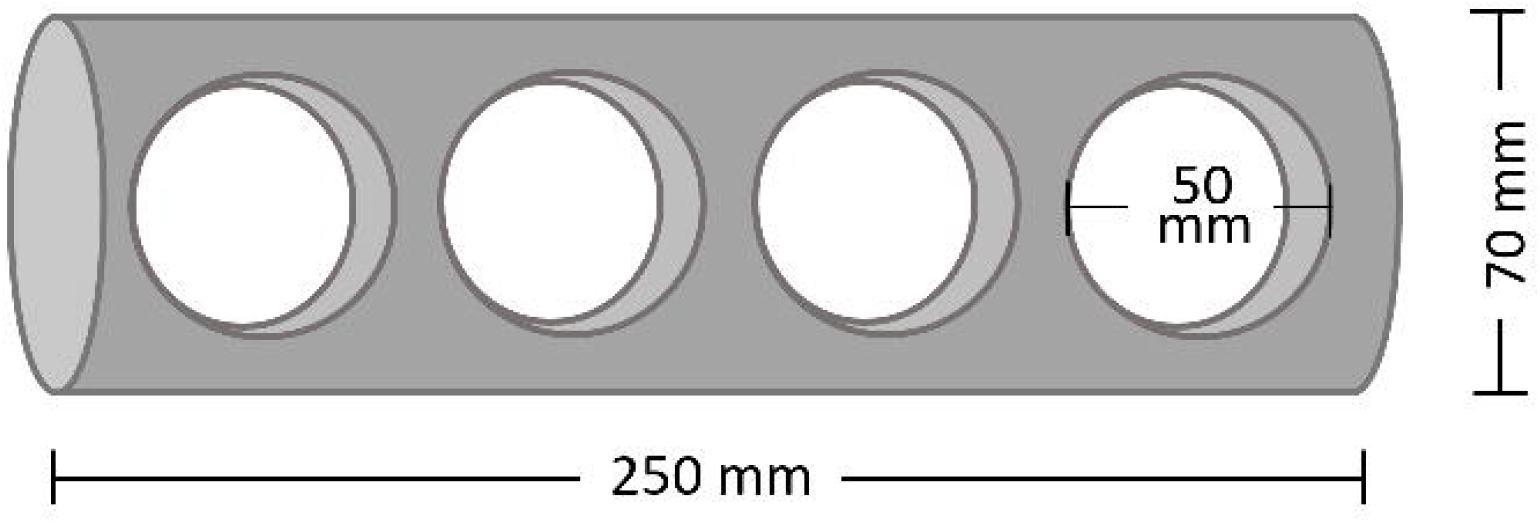
Representation of an empty ingrowth core seen from above/below. The cores were placed horizontally with the holes facing up and down, allowing roots to grow into the cores and through it

We applied the tracers Cs, Li, Se, Rb, and Sr by mixing them into the soil in the cores. During the first tracer round in 2016, Cs and Li were placed at 0.5 and 2.3 m depth respectively. During the 2^nd^ round Rb, Sr, and Se were placed at 0.5, 2.3 and 3.5 m depth respectively. In 2017 Cs, Li + Sr and Se + Rb was placed at 0.5, 2.3 and 2.9 m depth respectively (Table 3). Tracers where applied in the same concentrations as in the pilot experiment. However, in 2017, Sr application was doubled. For each core, 25 ml of the solution containing the tracer(s) in question was thoroughly mixed into the soil before filling it into the core.

**Table 3:**
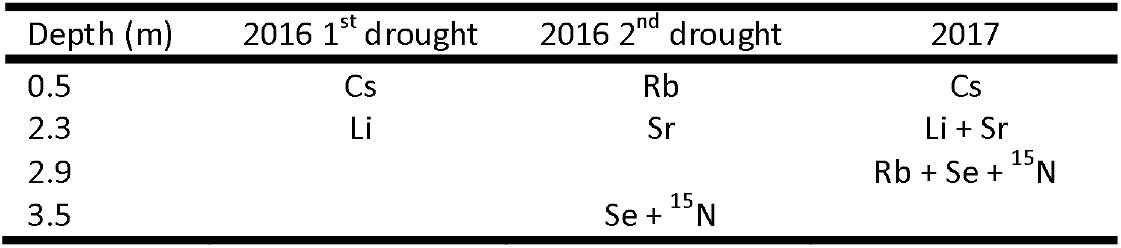
Depth of placement of each of the tracers during the l^5t^ and the 2^nd^ round in 2016 and in 2017.

We used a metal auger with the same diameter as the ingrowth cores to drill a hole, before placing the ingrowth core. We placed two ingrowth cores in each of the designated depths. The soil drilled out was stored frozen for later root washing and estimation of root length density (RLD), to give an indication of RLD prior to core insertion. For the 2^nd^ round in 2016 and the 2017 experiment, we reused as many holes as possible, therefore pre-samples do not exist for all treatments, and for some treatments only in fewer replicates. No pre-samples exist for the 1^st^ round for 0.5 m. Ingrowth cores were inserted 12 July and 23 August for the 1^st^ and 2^nd^ tracer round respectively in 2016 and removed again 28 July and 22 September respectively. In 2017, ingrowth cores were inserted 15 August and removed again 11 September (Table 2). The ingrowth cores were also stored frozen for later root washing.

We carefully washed the roots out of the ingrowth cores and the sampled bulk soil. We stored the roots in water at 5° C for up to 5 days until further measurement. To obtain RLD we placed the roots on a 200 x 250 mm tray and scanned using an Epson Perfection V700 PHOTO scanner on an 8-bit grayscale and 600 dpi. Root length measurements were obtained using WinRhizo^®^ software (Regent Instruments Canada Inc., 2006). The images were filtered using the WinRhizo setting of removing objects having a length?width ratio smaller than 4. In the setting of pixel classification, the automatic threshold was used.

We also used ^15^N as a tracer. We injected it into the soil volume at 3.5 and 2.9 m depth in 2016 and 2017 respectively. In 2016, we mixed 3 g Ca(^15^NO_3_)_2_ into 600 ml distilled water. Assuming that the soil contained 50 kg N ha^-1^ and that 10 % of the tracer would be taken up, this would result in a 2.3 times higher enrichment after application of tracer. In 2017, we used 2.82 g resulting in a calculated 1.65 times higher enrichment after application of tracer. We distributed the ^15^N solution among 100 sites, like this: We made two horizontal rows of each 10 equally distributed holes 5 cm above and below 2.3 m depth respectively. In each of these 20 holes, we injected 5 ml tracer distributed between five sites: 25, 20,15, 10 and 5 cm from the horizontal soil surface. We injected tracer 19 July in 2016 and 15 August in 2017.

### Biomass and tracer uptake

We harvested aboveground biomass 28 July and 22 September for the 1^st^ and 2^nd^ round respectively in 2016 and 11 September in 2017. We dried the biomass at 80°C for 48 hours. We also sampled and dried 2-3 leaves on 12 July 2016 and 14 August 2017 to be used as control samples for the tracer uptake. 1^st^ round biomass samples were used as a control for the 2^nd^ round in 2016, which was possible because a different set of tracers were used in the two rounds.

Samples were ground and analysed for tracer concentration. In 2016, the analyses of trace elements were made on an ICP-MS (Thermo-Fisher Scientific iCAP-Q; Thermo Fisher Scientific, Bremen, Germany) at University of Nottingham, School of Biosciences laboratory, and in 2017 they were analysed on an ICP-SFMS (ELEMENTXR, ThermoScientific, Bremen, Germany) at ALS lab, Luleå, Sweden. Analysis of ^15^N concentration was done by UC Davis stable isotope facility using a PDZ Europa ANCA-GSL elemental analyser interfaced to a PDZ Europa 20-20 isotope ratio mass spectrometer (Sercon Ltd., Cheshire, UK). Excess tracer concentration was calculated as in the pilot experiment

In order to identify whether tracer was present in a sample, we adapted the criteria proposed by Kulmatiski (2010) and modified by Beyer et al. (2016). If a sample had a concentration of a trace element at least 4 standard deviations (σ) higher than the control samples, tracer was assumed to be present.

### Statistics

In the pilot experiment, the effect of species (lucerne vs red beet) on aboveground biomass was tested in a one-way ANOVA, and the effect of soil depth and species on root intensity was tested in a mixed effects two-way ANOVA. Tube was included as a random effect to account for the fact that the different depths are not independent. We tested whether application of tracers resulted in a significant increase in uptake of each of the five trace elements from before to after application (Excess tracer concentration) in a threeway repeated measurements ANOVA with time (before/control & after tracer application), species and tracer type as factors. Furthermore, we tested whether excess tracer concentration differed among trace elements and species in a two-way ANOVA, using the control measurements as a baseline. Both approaches called for log transformation to meet the assumption of homoscedasticity, preventing us from testing both research questions in the same model. Se concentration was multiplied by 10^3^ as it was applied in a concentration of 10^-3^ compared to the other tracers.

In the rhizobox experiment the effect of treatment on aboveground biomass of chicory, black medic and ryegrass was tested in a mixed effects one-way ANOVA. Separate harvest of single plants allowed the inclusion of chamber as a random effect to account for the fact that the two intercropped species are not independent. The effect of treatment on RLD in the bulk soil and in the ingrowth cores was tested in a mixed effects one-way ANOVA, which was done because there were two ingrowth cores in each depth. We correlated RLD and the root intensities reported in Rasmussen et al. (2019a, preprint) obtained from observations of root development on the transparent surfaces of the rhizoboxes. We tested whether the correlation differed between bulk soil and ingrowth cores.

Test of significant tracer uptake and treatment effects on excess tracer concentration was tested as in the pilot experiment, with the exception that the tracers were tested in separate models. As for the biomass mixed effects models were used to account for single plant samples. We log-transformed whenever needed to meet the assumption of homoscedasticity. Effect of treatment on N concentration in chicory was tested in a mixed effects one-way ANOVA. We used separate models for each round and year in all cases. We correlated RLD and Li uptake from 2.3 m and tested the effect of treatment in a one-way ANOVA.

Differences were considered significant at P < 0.05. Data analyses were conducted in R version 3.4.4 (R Core team 2018). Tukey test P-values for pairwise comparisons were adjusted for multiplicity, by single step correction to control the family-wise error rate, using the multcomp package (Hothorn et al. 2008). Error bars represent 95 % confidence intervals.

## Results

### Pilot experiment

Both lucerne and red beet grew well. At harvest 1 September, 10 weeks after transplanting, lucerne aboveground biomass was 17.1 g and red beet biomass including the tuber was 25.8 g, which was significantly higher than lucerne biomass (Figure 2). Red beet had reached the bottom of the tube 20 July, just 4 weeks after transplanting, whereas lucerne roots were not observed in the bottom of the tubes before 3 August, 6 weeks after transplanting (data not shown). 10 August, 4 days after placing the pots with tracer below the tubes, root intensity was alike for the two species in the lower part of the tube. In the upper part of the tubes, red beet had more roots than lucerne (Figure 3).

**Fig. 2.**
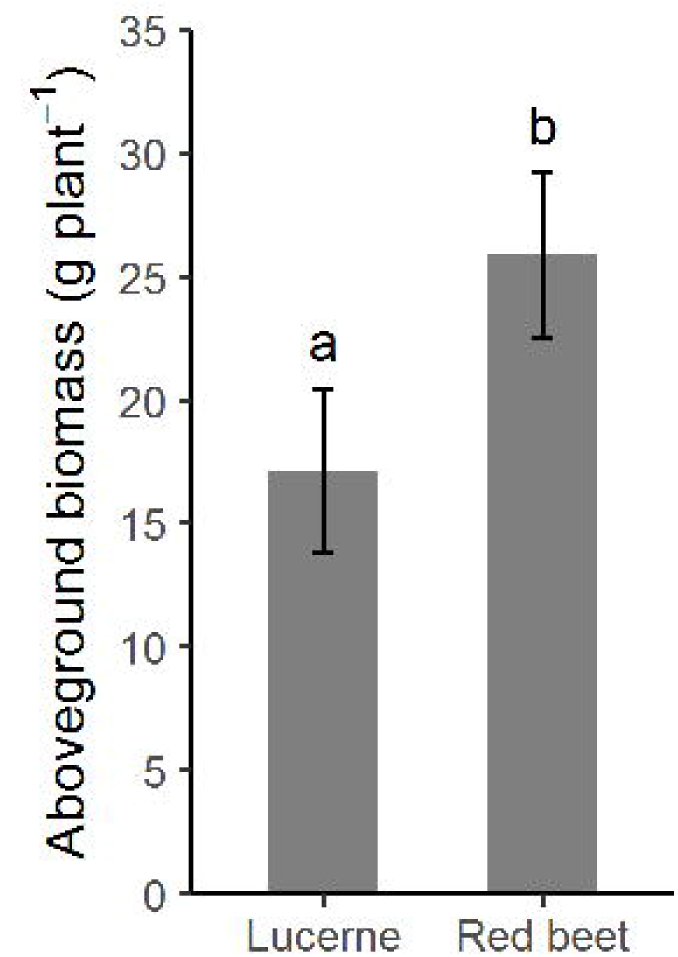
Biomass of lucerne and red beet harvested 1 September, 10 weeks after transplanting. Error bars denote 95 % confidence intervals and letters indicate significant differences

**Fig. 3.**
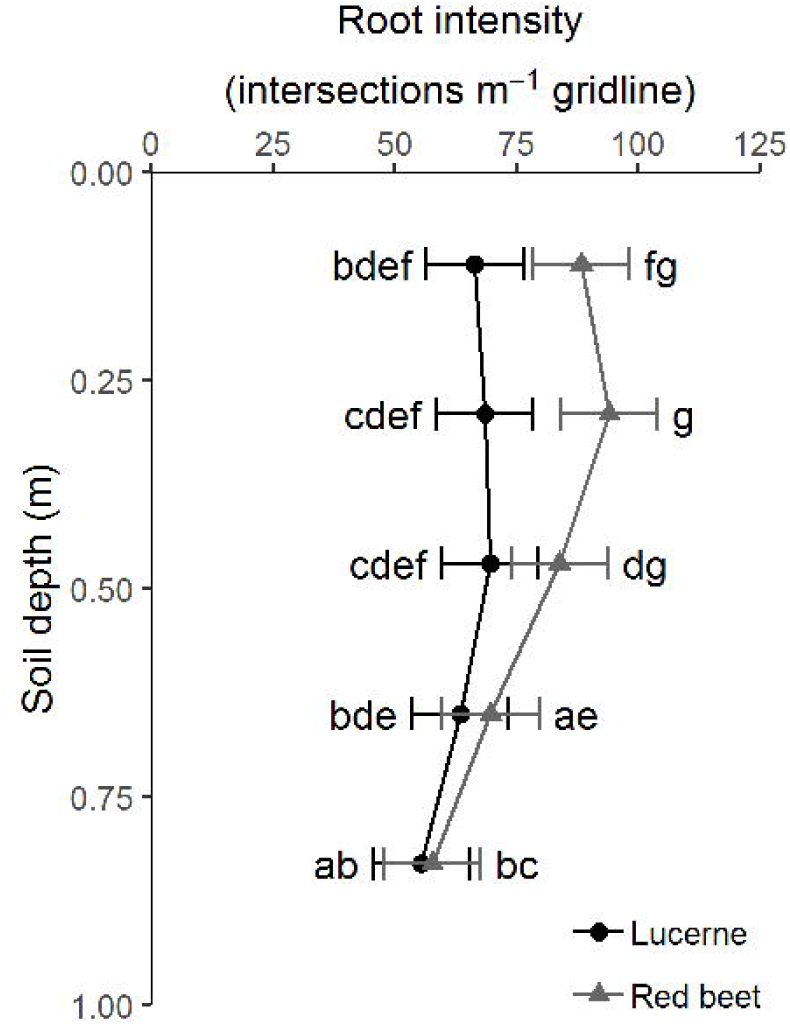
Root intensity of lucerne and red beet 10 August, 4 days after installing the pots with tracers and 7 weeks after transplanting. Error bars denote 95 % confidence intervals and letters indicate significant differences

Background concentration of the trace elements in lucerne was 6.6, 462, 0.22, 77, 0.13 μmol kg^-1^ for Li, Sr, Cs, Rb and Se respectively and likewise 6.5, 238, 0.19, 107, 0.12 μmol kg^-1^ red beet. The differences among species were not significant but among trace elements it was (data not shown). Tracer application increased the concentration of all the applied tracers in both species apart from Sr in lucerne. For lucerne, the increase was a factor 2.3,1.3, 793, 4.1, 37 for Li, Sr, Cs, Rb, and Se respectively and likewise 48, 3.2, 3244,16 and 35 for red beet. Differences in background levels among trace elements were not reflected in the levels of excess tracer concentration.

Red beet took up significantly more Li, Sr, Cs and Rb tracer than lucerne. The uptake of Se did not differ among the two species (Figure 4). For lucerne, the excess tracer concentration did not differ among Sr, Cs, and Rb. For Red beet, this was the case for Li, Sr, and Cs.

**Fig. 4.**
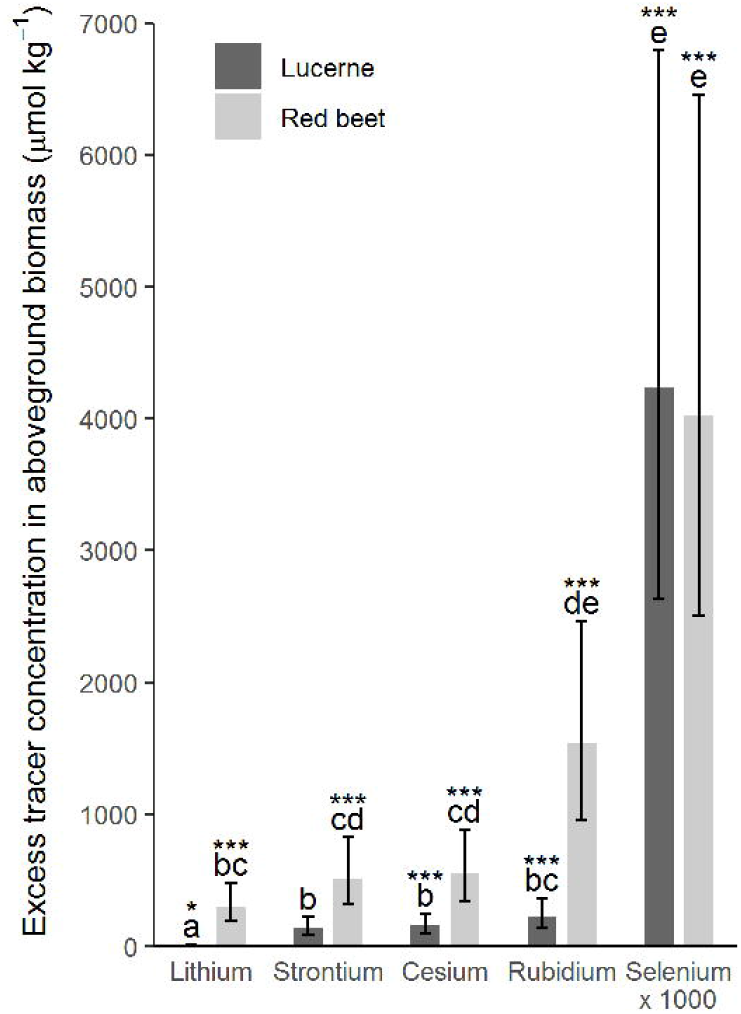
Excess tracer concentration of Cs, Li, Rb, Sr and Se x 1000 in aboveground biomass of lucerne and red beet after placing the tracers at 1 m depth. Error bars denote 95 % confidence intervals, letters indicate significant differences and *, ** and *** indicate that the excess tracer concentration is significantly different from 0 at P < 0.05, 0.01 and 0.001, respectively. Excess tracer concentration of Se has been multiplied by 1000 to normalize the amount of tracer applied among the tracers

### Main experiment

Plants grew well both years. In 2016, the chicory plants were in their second growth year and started flowering ultimo May, and in 2017 we used first-year chicory plants, which stayed in the vegetative state during the season. The three drought treatments lasted 31, 41 and 29 days, during which 135, 73 and 97 mm of water was excluded from the DS treatment in the 1^st^ and 2^nd^ round of 2016 and in 2017 respectively compared to the other treatments.

### Biomass and N content

We found no treatment effects on biomass in either of the rounds in 2016 (Figure 5). In the 1^st^ round, aboveground biomass harvested 28 July was 6.52 and 6.85 ton ha^-1^ in the WW and DS treatment respectively. In the 2^nd^ round harvested 22 September, plants were not allowed to regrow to their original size, and biomass was 2.51 and 2.03 ha^-1^ in the WW and the DS treatment respectively. In 2017, we harvested 11 September and chicory biomass was 4.65 and 3.64 ton ha^-1^ in the WW and DS treatment respectively, and 2.80 and 2.21 ton ha^-1^ when intercropped with black medic or ryegrass respectively. Biomass of black medic and ryegrass differed significantly and was 5.89 and 7.68 ton ha^-1^ respectively. Both intercropping treatments significantly reduced chicory biomass compared to the WW treatment (Figure 5). Biomass for the 1^st^ round 2016 and for 2017 has also been reported in Rasmussen et al. (2019a, preprint).

**Fig. 5.**
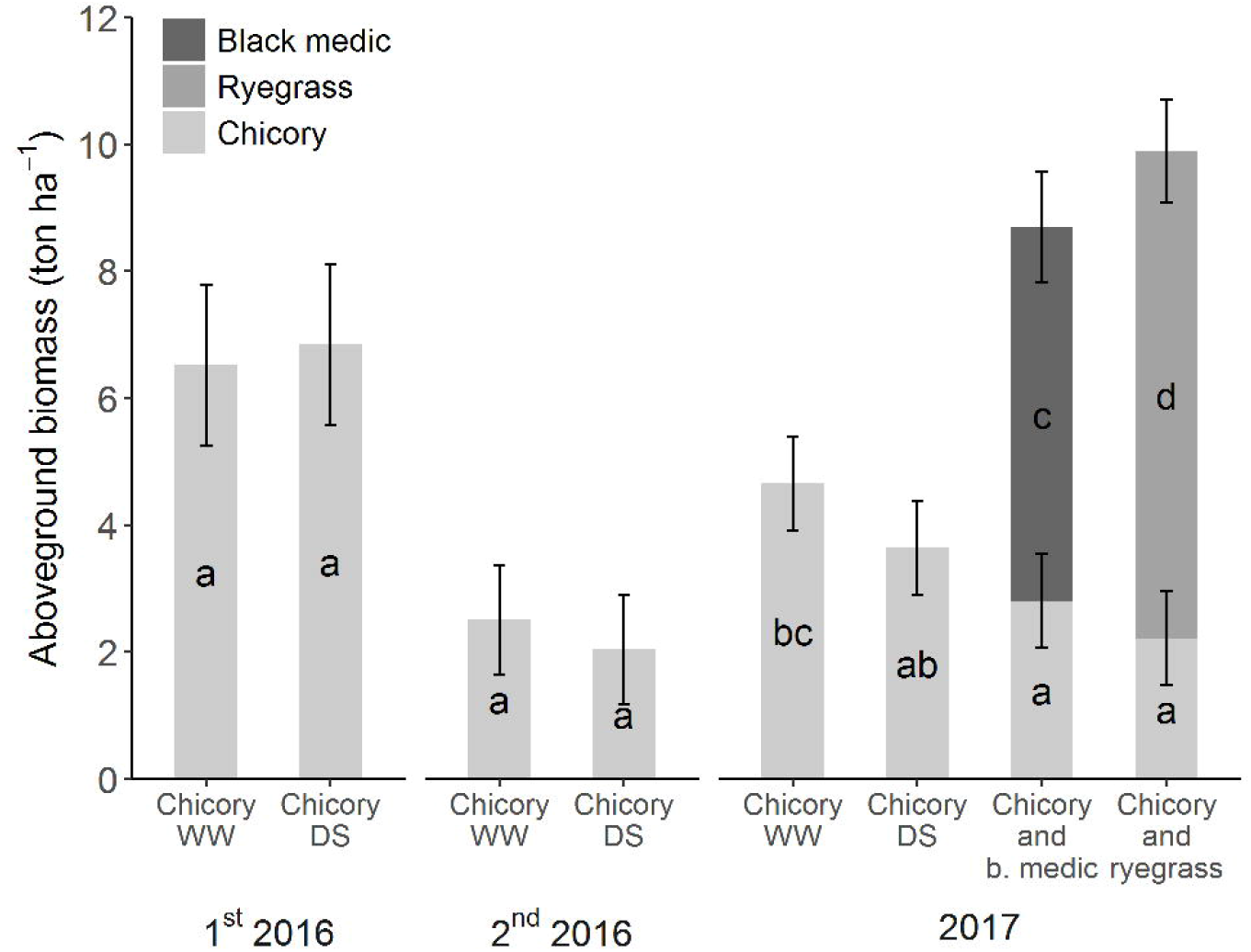
Biomass harvested 28 July and 22 September for the l^5t^ and 2^nd^ round in 2016 respectively and 11 September 2017. Error bars denote 95 % confidence intervals, and letters indicate significant differences. Each round and year was tested in separate models. Part of the data in the figure has also been shown in (Rasmussen et al. 2019b, preprint)

No treatment effects were seen on the N concentration in the plants in either of the rounds in 2016 or in 2017 (data not shown).

### Root length density

We correlated the RLD with the root intensities reported in Rasmussen et al. (2019a, preprint) obtained from observations of root development on the transparent surfaces of the rhizoboxes (Figure 6). We found that RLD in the bulk soil correlated well with root intensity (R^2^ = 0.5894). Roots grew into the ingrowth cores and there tended to be slightly higher RLD in the cores than in the bulk soil. The correlation with root intensity for RLD in ingrowth cores was weaker than for the bulk soil (R^2^ = 0.1622), due to increased variation.

**Fig. 6.**
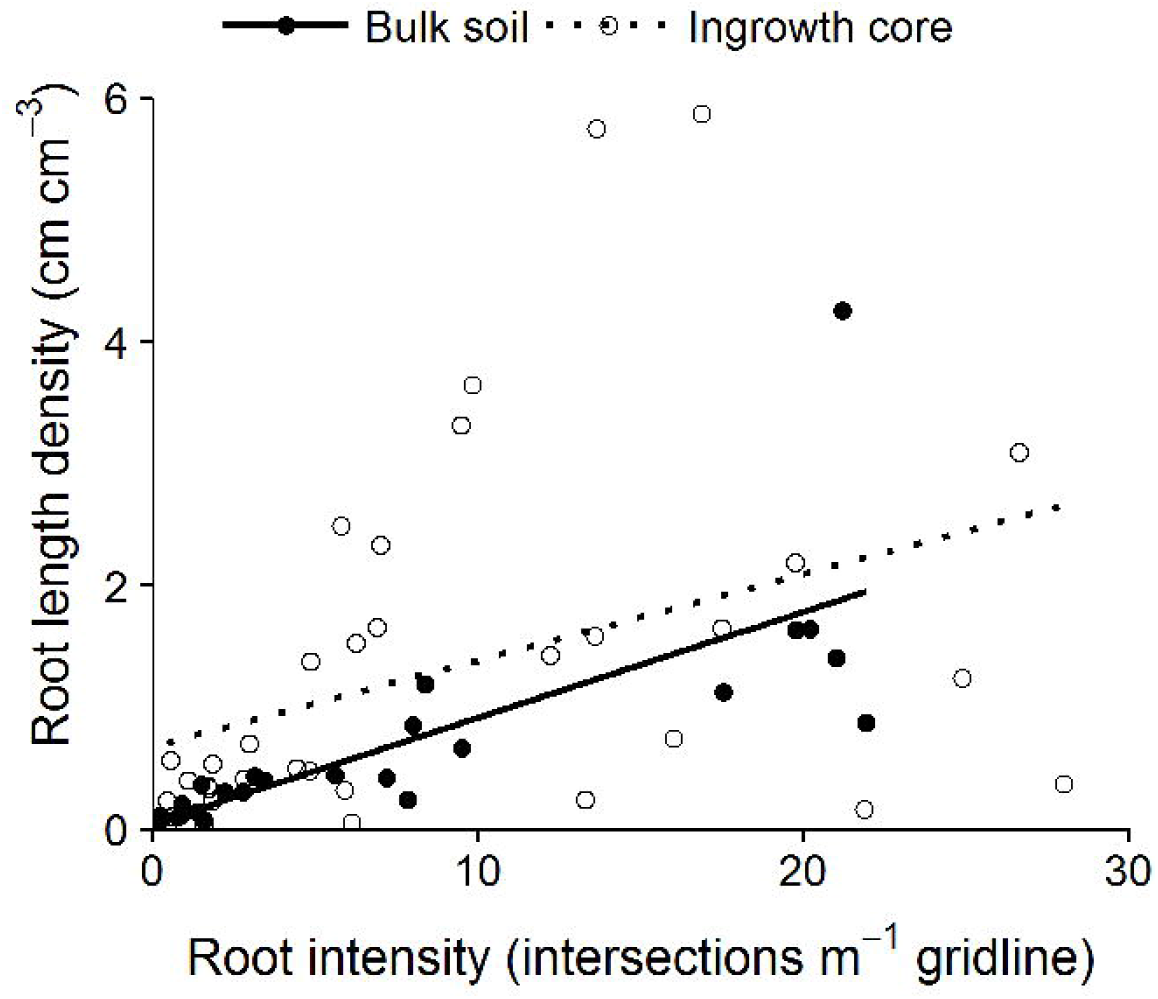
Regression between root intensity and root length density in bulk soil and ingrowth cores. R^2^ for the regressions is 0. 5894 and 0.1622 for bulk soil at ingrowth cores respectively

We did not find any treatment effect in the RLD in the 1^st^ round in 2016. In the 2^nd^ round in 2016, we found higher RLD in the DS than in the WW treatment in both 0.5 and 2.3 m depth, however only significant for the former. In 2017 we found a consistent tendency to a decreased RLD in the DS and the two intercropping treatments compared to WW at 2.3 m depth. This was also the case for 0.5 m depth, apart from the chicory and ryegrass having the most roots at this depth (Figure 7).

**Fig. 7.**
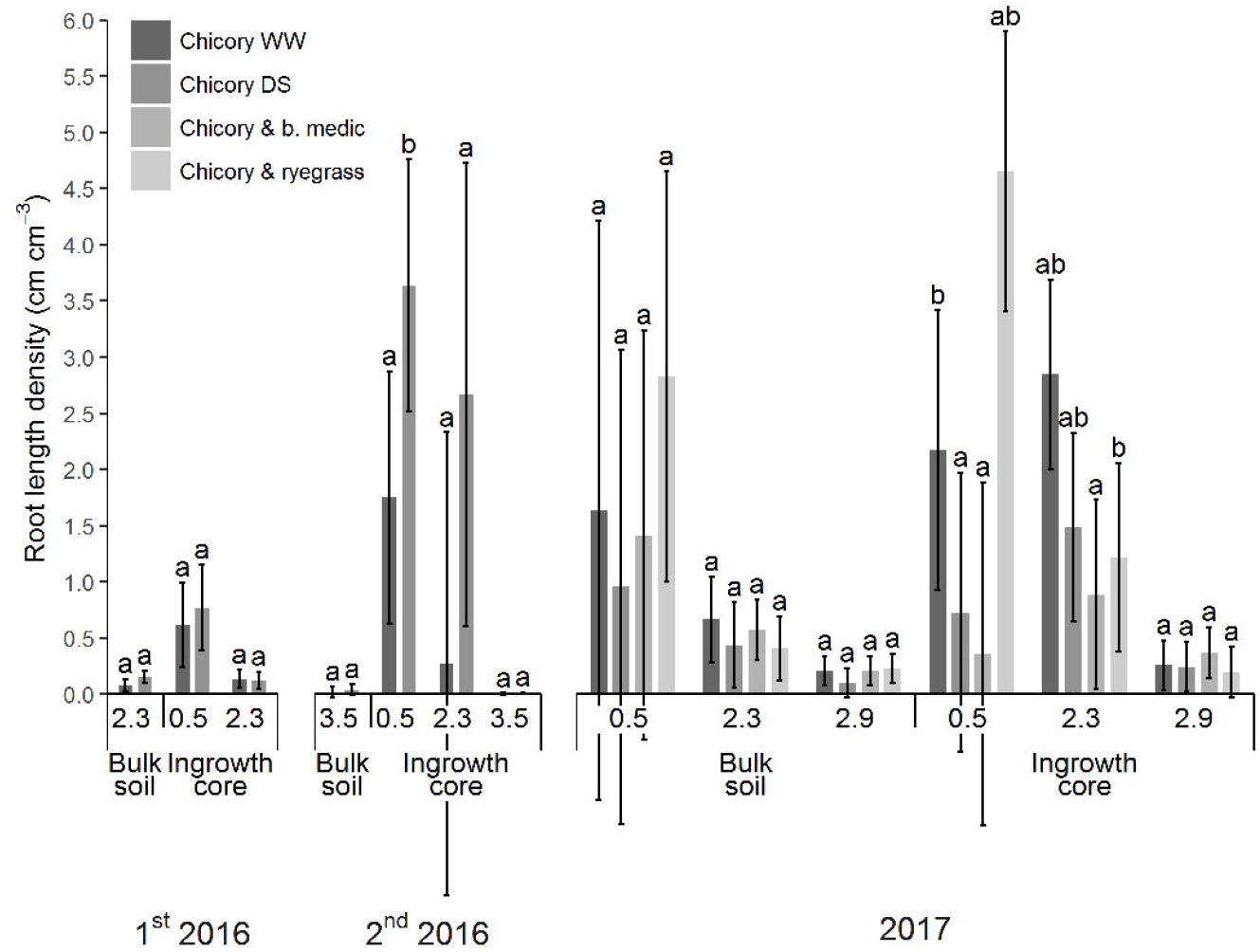
Root length density in the soil samples taken out when inserting the ingrowth cores (bulk soil) and inthe ingrowth cores, (ingrowth core). Ingrowth cores were inserted 12-28 July and 23 August to 22 September in l^5t^ and 2^nd^ tracer round in 2016 respectively. In 2017, ingrowth cores were inserted 15 August to 11 September. Error bars denote 95 % confidence intervals and letters indicate significant differences among treatments within round and depth

### Tracer uptake

In the 1^st^ round in 2016, we found a significant uptake of the Li tracer, which had been injected at 2.3 m depth but not of Cs which had been injected at 0.5 m depth. In the second round, we found a significant uptake of the Rb tracer injected at 0.5 m and of Sr injected into 2.3 m depth, but only in the wet treatment. For the tracers injected into 2.3 and 3.5 m depth, there was a tendency to a higher excess tracer concentration in WW chicory than in DS chicory, but the difference was only significant for Sr taken up from 2. 3 m (Figure 8).

**Fig. 8.**
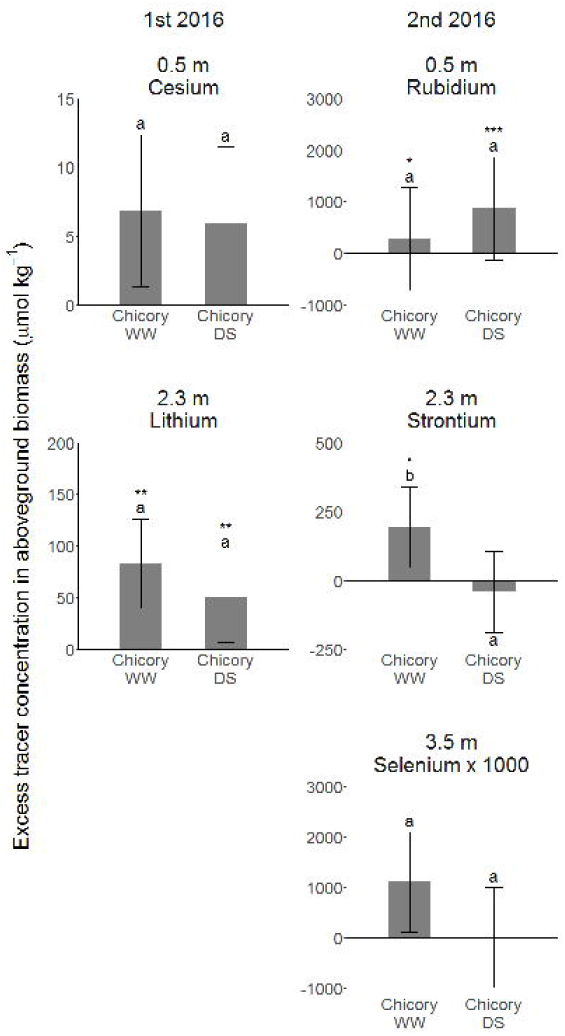
Excess tracer concentration in aboveground biomass after injection of Cs at 0.5 m and Li at 2.3 m depth at the l^st^ round in 2016 and injection of Rb at 0.5 m, Sr at 2.3 m and Se at 3.5 m depth at the 2^nd^ round in 2016. Error bars denote 95 % confidence intervals, letters indicate significant differences and *, **and *** indicate that the excess tracer concentration is significantly different from 0 at P < 0.05, 0.01 and 0.001, respectively. Excess tracer concentration of Se has been multiplied by 1000 to normalize the amount of tracer applied among the tracers

In 2017, we only found that tracer application significantly increased uptake of Cs from 0.5 in WW chicory and in ryegrass intercropped with chicory and of Li from 2.3 m depth in WW chicory. Excess tracer concentration of Cs was also significantly higher in WW chicory and ryegrass intercropped with chicory than in black medic or chicory in the other treatments. Excess tracer concentration of Li was higher in WW chicory, chicory intercropped with black medic or ryegrass than in black medic intercropped with chicory. Whereas we did not find tracer uptake to be significant for Rb injected into 2.9 m depth we did find significant treatment effects, as chicory intercropped with black medic had a higher excess tracer concentration than black medic intercropped with chicory. Though not significant, we did find a general pattern of the tracers taken up from 2.3 and 2.9 m depth, as excess tracer concentration was higher in the WW chicory and chicory intercropped with black medic or ryegrass than in the other chicory treatments (Figure 9).

**Fig. 9.**
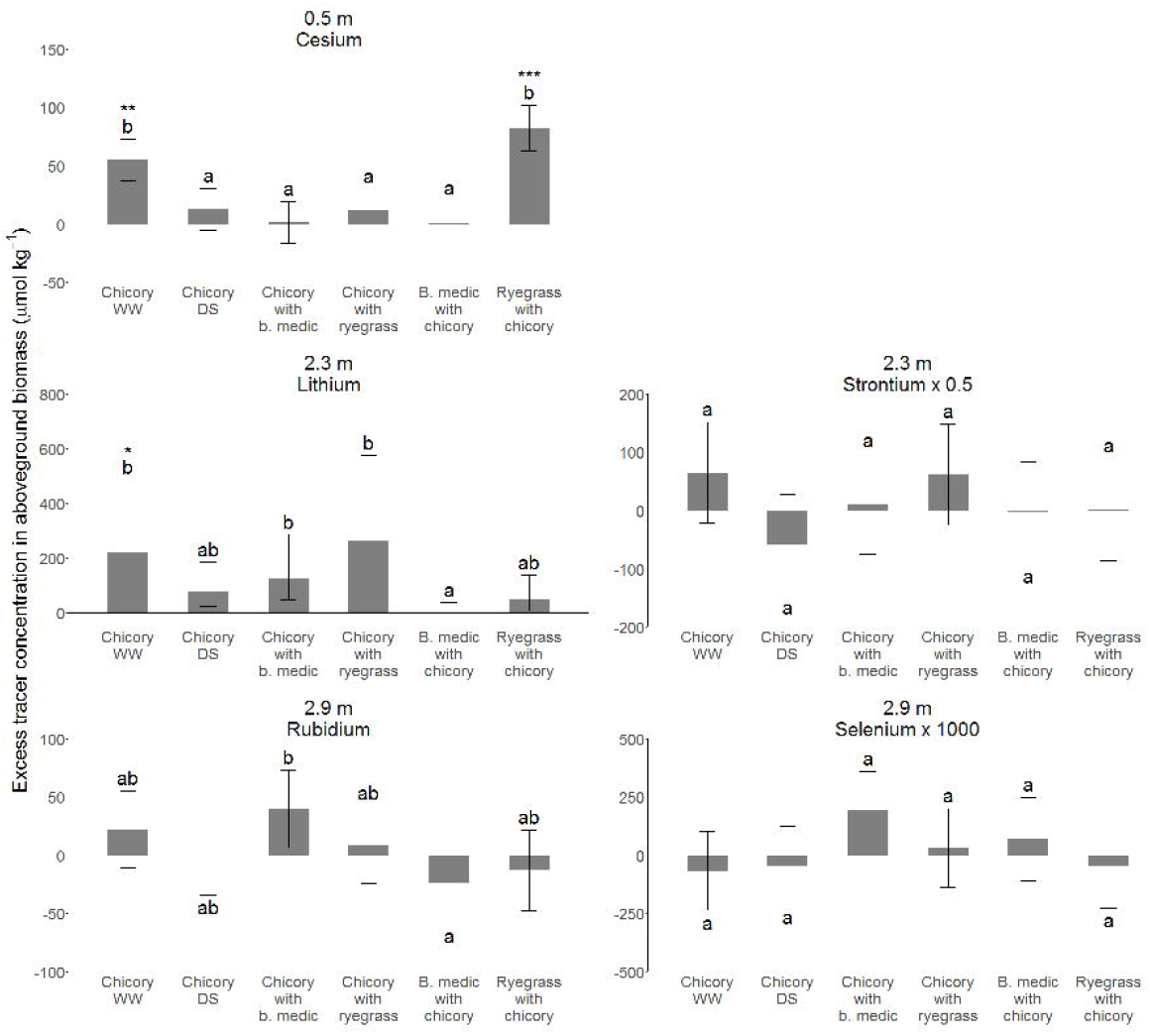
Excess tracer concentration in aboveground biomass after injection of Cs at 0.5 m depth, Li and Sr at 2.3 m depth and Rb and Se at 2.9 m depth in 2017. Error bars denote 95 % confidence intervals, letters indicate significant differences and *, ** and *** indicate that the excess tracer concentration is significantly different from 0 at P < 0.05, 0.01 and 0.001, respectively. Excess tracer concentration of Sr has been multiplied by 0.5 and Se by 1000 to normalize the amount of tracer applied among the tracers

We found a tendency to a steeper increase in excess Li tracer concentration in the biomass per root length in chicory DS and chicory intercropped with ryegrass than chicory in the two other treatments in 2017, indicating a more effective Li uptake from 2.3 m depth in these treatments. The differences were not significant though excess tracer concentration per root length density was 3 and 6 times higher respectively. We did not find the same tendency in the 1^st^ round in 2016. Similar correlations could not be carried out for other tracers injected into either 2.3 or 3.5 m depth, due to too many samples without tracer uptake (Figure 10).

**Fig. 10.**
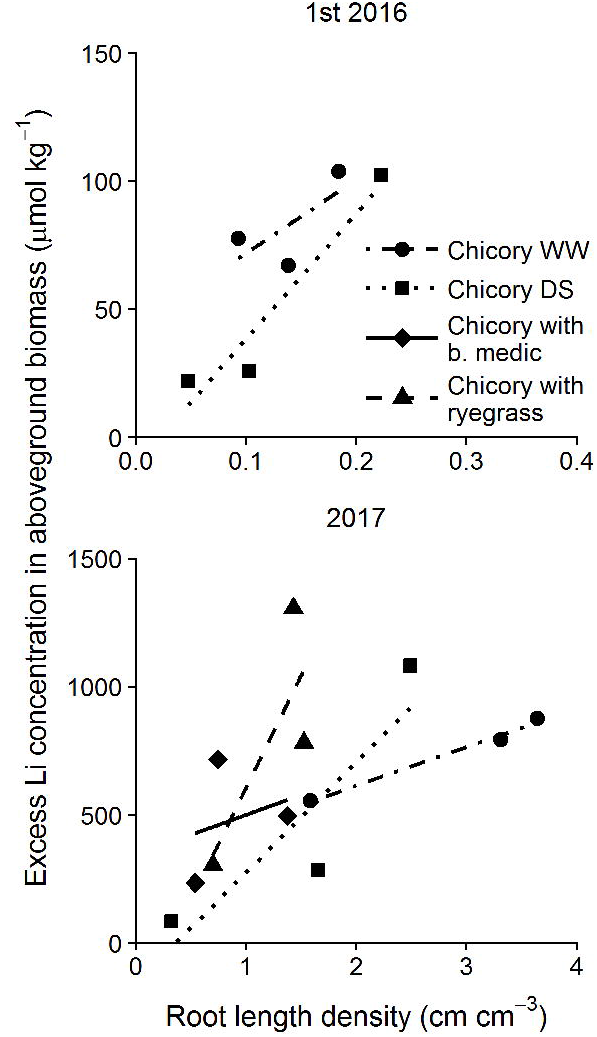
Regression between root length density in the ingrowth cores at 2.3 m depth and excess tracer concentration of Li in aboveground biomass in the l^5t^ round in 2016 and in 2017. As black medic and ryegrass are shallow rooted, the roots are assumed to originate from chicory only. Note the factor 10 between the axes of the two plots

By comparing the concentration of the five trace elements in the control samples taken before applying tracers in 2017, we could test whether tracer application in 2016 had contaminated the chambers as three of the chambers had not been used for tracer application in 2016. We found that the background concentration of Li, Sr, and Cs was higher in the chambers used in 2016 than those only included in 2017, however only significant for Cs (Figure 11). ^15^N had been used in all chambers in 2016 and the ^15^N concentration in the control samples increased significantly from 22.5 mg kg^-1^ (95 % confidence interval 8.2 – 36.7) in 2016 to 64.9 mg kg^-1^ (56.6 – 71.1) in 2017.

**Fig. 11.**
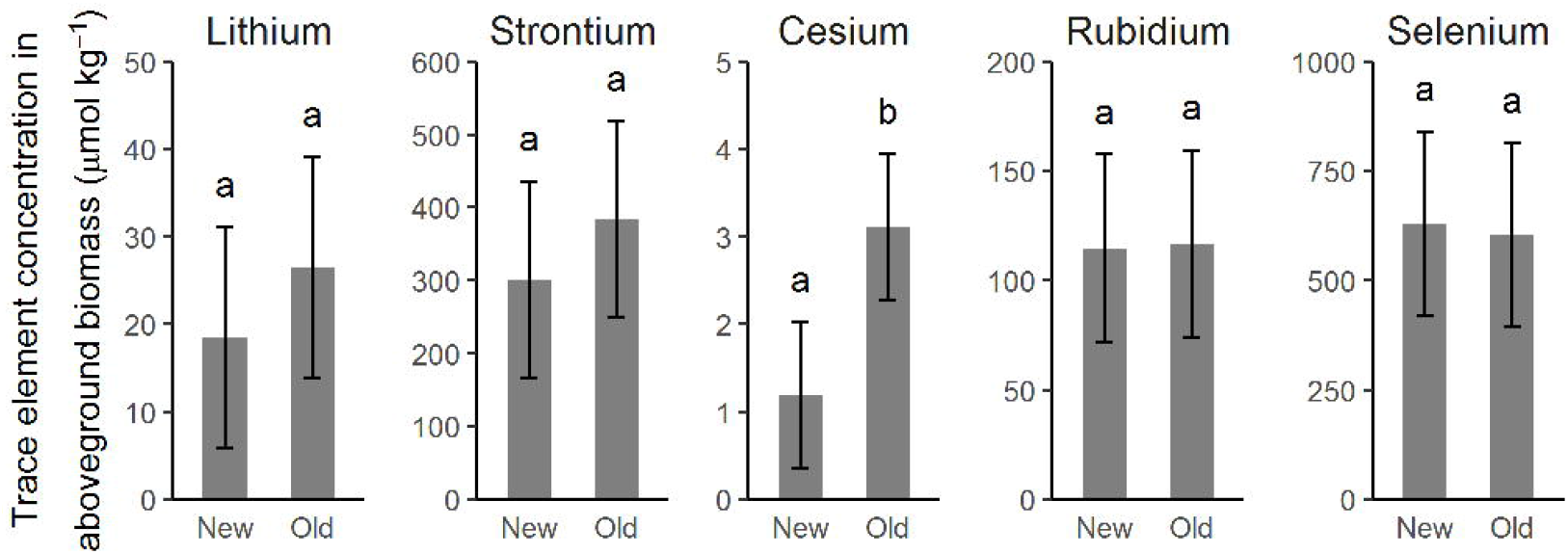
Concentration of the trace elements Li, Sr, Cs, Rb, and Se in aboveground biomass before applying tracers in 2017. New chambers denote plants grown in chambers where tracers have never been applied, and reused chambers denote plants grown in chambers receiving tracers in 2016. Error bars denote 95 % confidence intervals and letters indicate significant differences between plants grown in new and reused chambers for each trace element

Placing the trace element tracers in two ingrowth cores per depth limited the number of plants accessing the tracers. In the 1^st^ round in 2016 and in 2017 we found that 40-75 % of the chicory plants exposed to mean of the control samples, which according to our criteria indicated that these plants had taken up tracer (Figure 12 and 13). This, however, did not apply to Sr in 2017, which was not taken up by any of the chicory plants. In the 2^nd^ round in 2016 no chicory plants took up tracer from 0.5 m depth and only 25 % of the chicory plants from 2.3 m. 13 % of the chicory plants took up tracer from 3.5 m depth in the 2^nd^ round in 2016 and 3 % from 2.9 m depth in 2017.

**Fig. 12.**
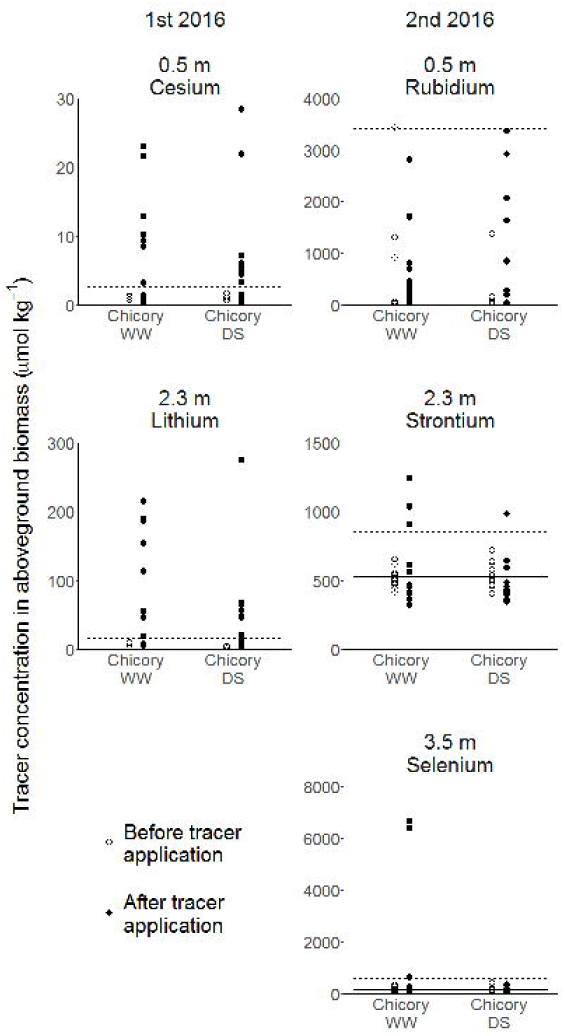
Trace element concentration in single plant aboveground biomass before and after injection of Cs at 0.5 m and Li at 2.3 m depth at the l^5t^ round in 2016 and injection of Rb at 0.5 m, Sr at 2.3 m and Se at 3.5 m depth at the 2^nd^ round in 2016. Solid and dashed lines denote mean and 4 standard deviations (σ) above mean concentration before tracer application. n=3 fora l^5t^ round before tracer application and n=12 for the rest

**Fig 13.**
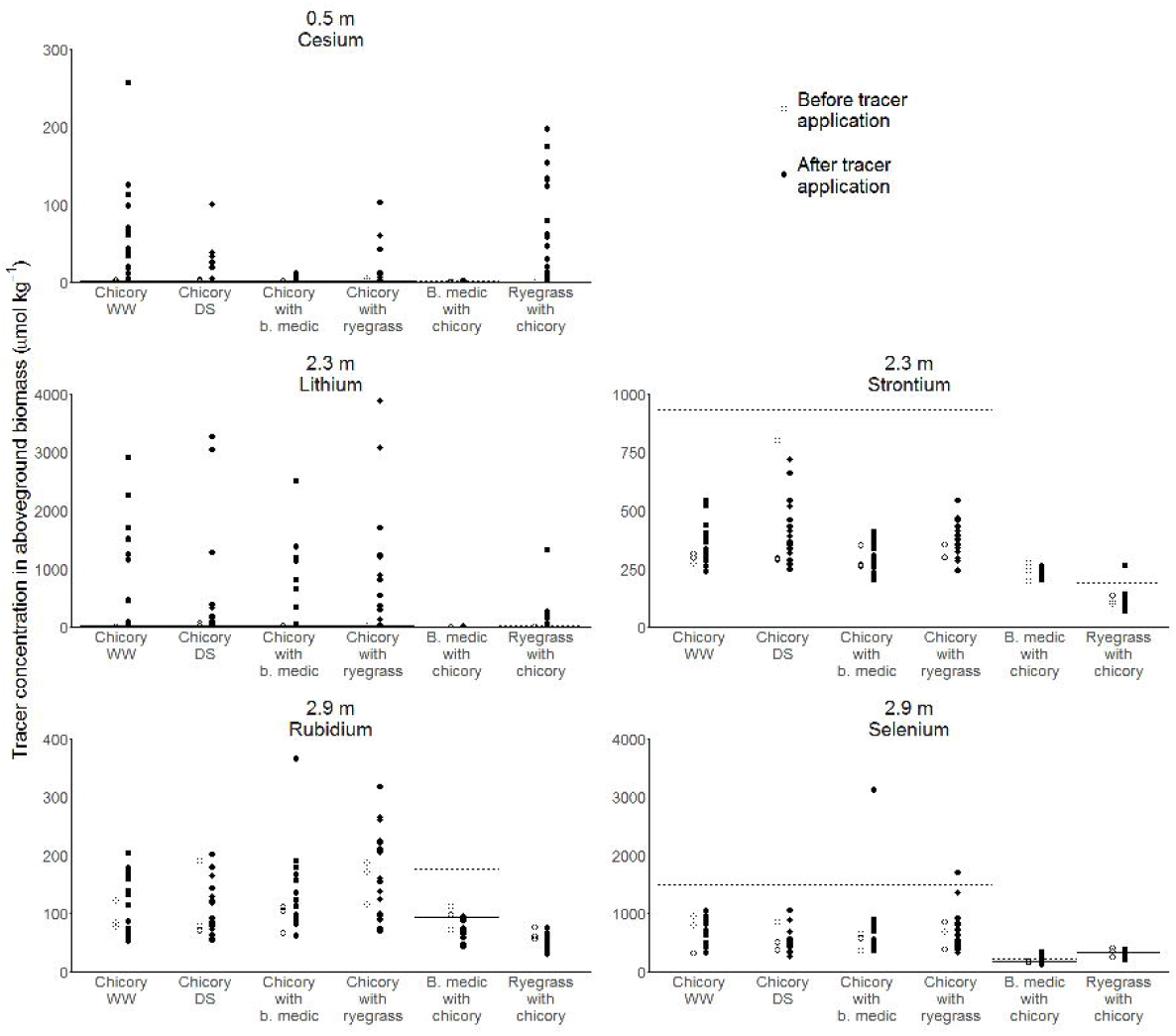
Trace element concentration in single plant aboveground biomass after injection of Cs at 0.5 m depth, Li and Sr at 2.3 m depth and Rb and Se at 2.9 m depth in 2017. Solid and dashed lines denote mean and 4 standard deviations (σ) above mean concentration before tracer application for each species. n=3 for before tracer application and n=18 for chicory and n=15 for black medic and ryegrass after tracer application

All the ryegrass plants and 7 % of the black medic plants took up tracer from 0.5 m depth in 2017. From 2.3 m depth, 53 % and 7 % of the ryegrass plants took up Li and Sr respectively, whereas this only applied to 7 and 0 % of the black medic plants. Ryegrass did not access tracer from 2.9 m, but 67% of the misleading.

^15^N concentration in the biomass significantly increased after tracer application in 2016, but not in 2017. Excess tracer concentration was decreased by 50 % in the DS treatment compared to the WW treatment in 2016, but the difference was not significant. In 2017, excess tracer concentration was significantly higher in chicory than in ryegrass when the two species were intercropped (Figure 14).

**Fig 14.**
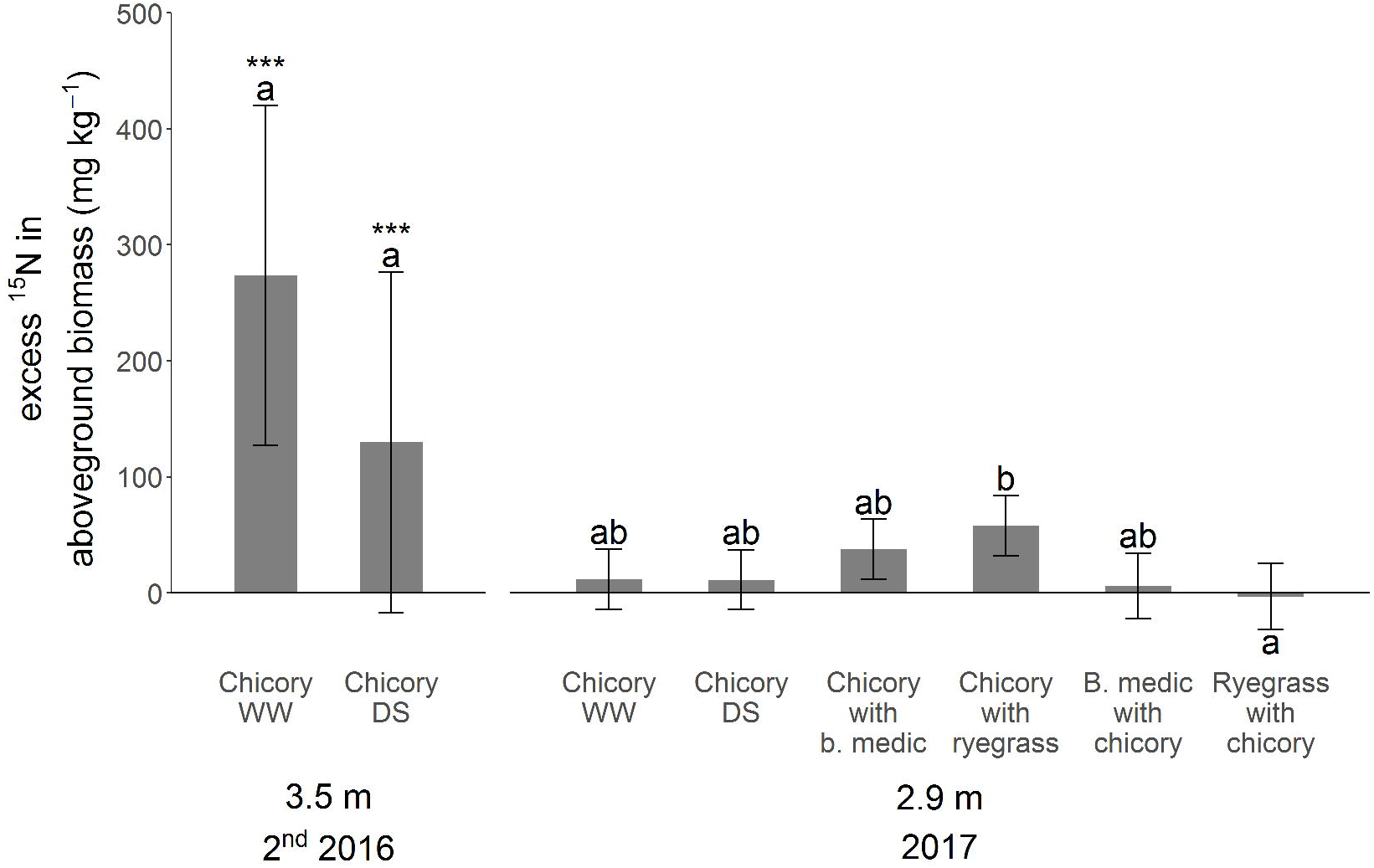
^15^N concentration aboveground biomass after injection at 3.5 m depth at the 2^nd^ round in 2016 and 2.9 m depth in 2017. Error bars denote 95 % confidence intervals, letters indicate significant differences and *, ** and *** indicate that the excess tracer concentration is significantly different from 0 at P < 0.05,0.01 and 0.001, respectively. Each year was tested in separate models

According to our criteria, all chicory plants took up tracer from 3.5 m depth in 2016, and 41 % in 2017. Also, 13 % of the black medic and the ryegrass plants took up tracer. The fact that variation among control samples was very small these numbers might be misleading (Figure 15). A visual evaluation suggests that 5 chicory plants took up tracer in 2016 and 1-3 in 2017, as these are showing ^15^N concentrations substantially higher than the other plants.

**Fig 15.**
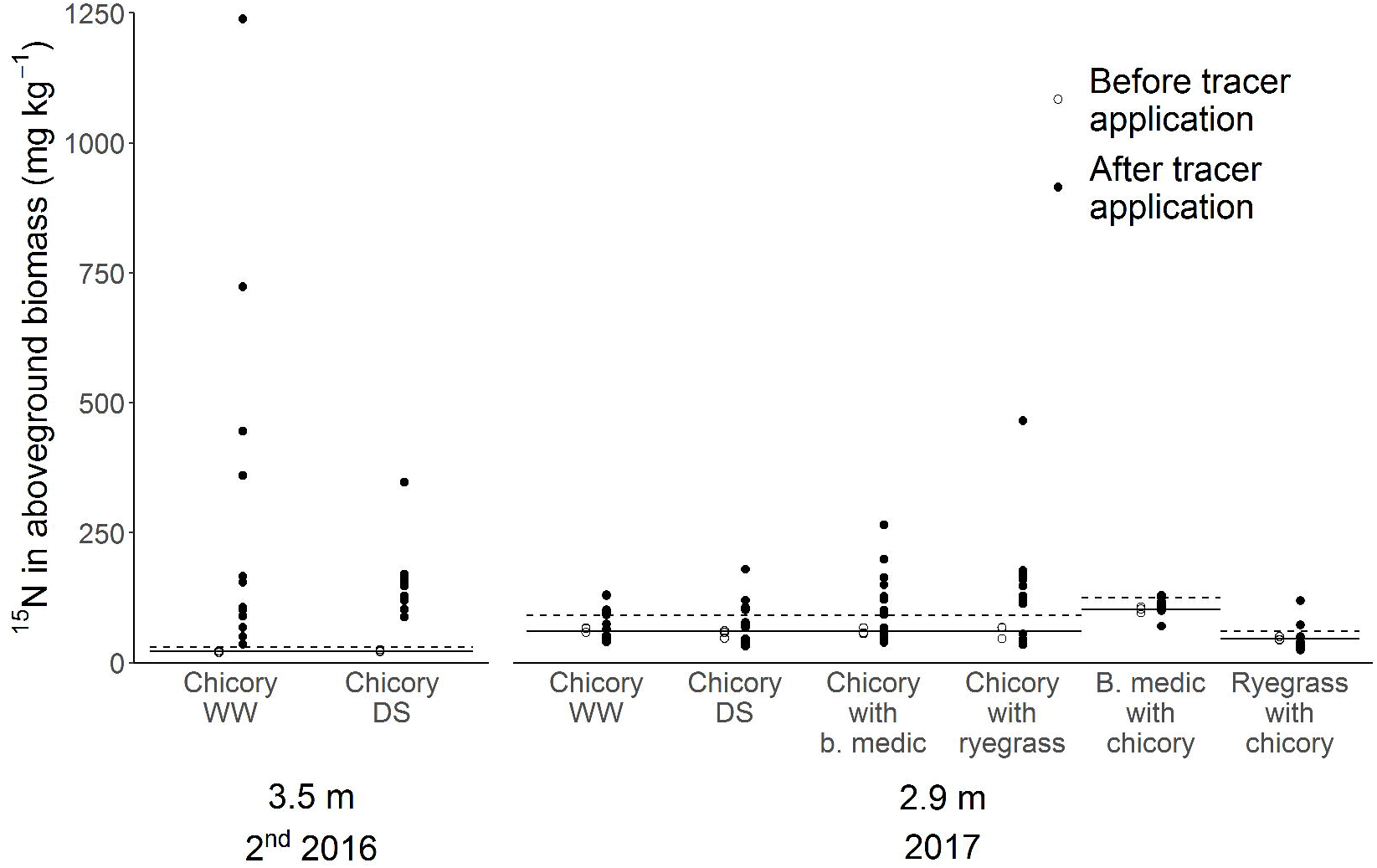
^15^N concentration in single plant aboveground biomass after injection at 3.5 m depth at the 2^nd^ round in 2016 and 2.9 m application. n=3 for before tracer application in the 2^nd^ round in 2016 and n=12 after tracer application. In 2017 n=3 for before tracer application and n=18 for chicory and n=15 for black medic and ryegrass after tracer application

## Discussion

### Comparability among trace element tracers

Our pilot study confirmed that applying the trace elements Cs, Li, Rb, Sr, and Se could trace root activity at 1 m depth. For the less mobile trace elements, excess tracer concentration was higher in red beet than in lucerne, whereas no differences were seen for Se which has a higher soil mobility. As the background concentrations of the trace elements did not vary between the two species, the difference does not seem to originate from variation in affinity. Red beet had more roots in the profile at tracer injection, suggesting that it was able to grow more roots into the pot with tracers, explaining the higher uptake of Cs, Li, Rb and Sr in red beet than in lucerne. The mobile Se can probably be exploited to the same extent by both species despite differences in root density. Contrary to our findings, others have found differences in tracer affinity among species (Tofinga and Snaydon 1992; Hoekstra et al. 2014).

Application of equimolar amounts of the four less mobile trace elements resulted in similar excess tracer concentrations within species. Thus, data from our pilot study suggest that the trace elements can substitute each other in multiple tracer studies. In such studies, different trace elements representing the same nutrient or simply representing uptake activity are injected into different soil depths within the same experimental units. However, Sr and Li excess tracer levels were not comparable in our main experiment in 2017 despite that they were both mixed into the same ingrowth cores at 2.3 m. Though some studies have found strong correlations between uptake of various pairs of trace element tracers, the number of studies examining the relationships are still limited and discrepancies exist regarding which tracer pairs correlate well and under which circumstances. Hoekstra et al. (2014) found a strong correlation between plant excess tracer concentrations of Li, Rb, and Cs, but found that the correlation was affected by a drought treatment and that a correction factor between pairs of tracers is required for quantitative comparison. Collander (1941) too, found that the uptake Rb and Cs was very similar by a range of plant species from nutrient solutions and Gockele et al. (2014) confirmed the strong correlation between uptake of Li and Rb. Mamolos et al. (1995) on the other hand found that the correlation between Cs and Li uptake was only significant in some of the tested cases, but found a strong correlation between Sr and Cs uptake. Both Gockele et al. (2014) and Mamolos et al. (1995) found a weak correlation between the uptake of Sr and Li. These findings concurrently emphasize the prospects of using the trace elements Cs, Li, Rb, and Sr as multiple tracers within experimental units to reduce between-plot variation. However, they also call attention to possible limitations of direct quantitative comparisons, and especially in comparisons among species. The correlations might be affected by the concentration of other nutrients such as potassium and calcium (Cline and Hungate 1960; Ozaki et al. 2005; Kobayashi et al. 2016)

### Deep root activity

Our results demonstrated that chicory acquired trace element tracers applied to 2.3 m depth and N from 3.5 m. We also found indications of trace element uptake from 2.9 and 3.5 m. Contrary to our expectations we found an overall tendency to a lower excess tracer concentration from all depths in chicory exposed to drought than in well-watered chicory, however only significant for Sr taken up from 2.3 m in 2016. I.e. even where drought tended to decrease biomass of chicory, nutrient uptake was decreased even more than the biomass and no compensatory uptake from deeper soil layers was observed. Hoekstra et al. (2015) found that the proportional Rb and ^15^N uptake from 0.35 m depth compared to 0.05 m depth in chicory increased as a response to drought, but this was mainly due to a decrease in uptake from 0.05 m depth during drought. They also found that during drought chicory had a higher proportional uptake of both tracers from 0.35 m soil depth than ryegrass, and this actually originated from a different amount of water taken up at 0.35 m depth. However, it could be caused by differences in the depth of drought, as the two species might not have had the same water use, thus we question whether this higher proportional uptake can be explained by differences in rooting depth. Our results indicate that both species have dense root systems still at 0.5 m depth and that even intercropped ryegrass has more roots that sole cropped chicory at this depth.

Our results concurrently gave indications of a compensatory tracer uptake from 2.3 and 2.9 m depth in intercropped chicory compared to well-watered chicory, which is supported by the fact that chicory plants only showed tracer concentrations higher than 4σ above mean of control when intercropped. In addition, ^15^N tracer concentration in chicory intercropped with ryegrass was significantly higher than in ryegrass, though the tracer uptake in itself was not significant. Whereas the drought was imposed during the season, chicory was influenced by intercropping from the beginning, which might explain why we observe an effect of intercropping, but not of drought.

Though we did not observe increased deep nutrient uptake in absolute terms as a response to the drought we did find a relative increase, as nutrient uptake per root length increased at 2.3 m depth in 2017, though not significant. This was also the case for the chicory when intercropped with ryegrass. Ryegrass is unlikely to reach 2.3 m depth in the field (Thorup-Kristensen 2006), but the fact that half of the ryegrass repacked soil. Thus, as some of the roots in the cores at 2.3 m depth must have contained ryegrass roots, the relationship between chicory Li uptake and RLD at 2.3 m depth was even steeper than the data shows. In a study of eucalyptus trees, da Silva et al. (2011b) likewise found that uptake rate per unit of fine root length density increased with depth down to 3 m, to some extent counterbalancing the lower root length density in deeper soil layers.

In the 1^st^ round in 2016, the drought treatment had little impact on plant behavior. RLD in the bulk soil and in the ingrowth cores did not differ among the two treatments, neither did root intensity estimated from root growth on the soil-rhizotron interface (Further details in Rasmussen et al. 2019a, preprint), aboveground biomass, N concentration in aboveground biomass nor tracer uptake from 0.5 and 2.3 m. Though water uptake was impaired at 0. 5 m (Rasmussen et al. 2019a, preprint), this was not reflected in the nutrient uptake. The drought might have been too short to affect biomass and nutrient uptake.

In the 2^nd^ round, biomass reduced more than 50 % compared to the first round because the plants did not have time to regrow to the same size after being cut at ground-level. Still, root growth into the ingrowth cores was more than 3 times as high as at 0.5 m depth, and in the dry treatment also at 2.3 m depth compared to the 1^st^ round. RLD in the ingrowth cores were higher in chicory exposed to drought than when well-watered in both 0.5 and 2.3 m, but the treatments did not differ in either aboveground biomass or N concentration in aboveground biomass. This result was puzzling, especially because the opposite was observed in 2017. We suggest that the higher root growth in the drought-stressed chicory could be a result of these plants being in the well-watered treatment before cutting, as we swopped the chambers used for the two treatments after the 1st round. The prior well-watered plants could thus have been stronger and have a head start, offsetting the 2^nd^ round drought stress. Despite the higher RLD in the drought-stressed chicory at 2.3 m we found a significant higher Sr excess tracer concentration in the well-watered chicory and a tendency to a higher Se and ^15^N uptake from 3.5 m depth.

### Reflections on methodology

We placed the trace element tracers in ingrowth cores to avoid contamination of the soil in the facility. Still, we did find a tendency to contamination, which could potentially increase over time by repeated use of the same tracers. The contamination could originate from either tracer leaching from the ingrowth cores into the bulk soil or turnover of roots containing tracers from first years experiment. Sayre and Morris (1940) did not find signs of Li leaching during a growing season. Others found some leaching but never more than 10 cm (Pinkerton and Simpson 1979; Hoekstra et al. 2014; Gockele et al. 2014), which is of little importance for tracer experiment like ours with large distances between tracer application depths.

The use of ingrowth cores gave a patchy distribution of the tracers, which especially in the deeper application sites, critically limited the number of plants accessing the tracer. Comparing the number of plants taking up Se and ^15^N tracer from 3.5 m depth in the 2^nd^ round in 2016 shows that a more evenly distribution of tracer does increase the number on plants getting in contact with the tracer. This would call for placing more ingrowth cores with the current concentration of the tracers rather than increasing the concentration to detect differences in deep nutrient uptake among treatments. However, the higher number of plants containing Li than Sr tracer in 2017 indicates that the Sr tracer concentration was too low, as the tracers are taken up from the exact same ingrowth cores inserted into 2.3 m depth. This was the case despite that we had doubled the Sr application rate in 2017, and that we found similar uptake rates of Li and Sr in our pilot study. Placing more ingrowth cores calls for drilling the holes, and placing ingrowth cores dummies before the roots are growing into the soil, to avoid disturbance of the root system when inserting ingrowth cores. Hoekstra et al. (2014) found that even increasing the number of injection sites from 36 to 144 rrf^2^ of Li injection into 5 cm depth tended to decrease variability of tracer uptake, but such high number of sites are not compatible with ingrowth cores. Fox and Lipps (1964) found that tracer uptake was lower but so was the variability when increasing the number of injection sites from 1 to 4 holes 0.3 m^2^.

We found a tendency to a higher RLD in the cores than in the bulk soil, which is likely caused by the nitrogen hotspots created by the application of NO_3_ based tracers in the cores. The variation in RLD also tended to be higher in the cores than in the bulk soil, especially in layers with low root intensity, which was primarily in the deeper soil layers. So what we found was not just that far from all plants reached an ingrowth core in the deeper soil layers, but also that not all ingrowth cores in the selected depths were colonized by roots. The fact that RLD in the cores is continuously increasing and that the age of the roots in the cores does not reflect the age of the roots in the bulk soil prevents direct comparisons in nutrient uptake from the cores and the bulk soil.

We speculate that placing the holes of the ingrowth cores facing towards the sides instead of up and down or choosing a design with access on a larger part of the ingrowth cores surface could have increased the number of colonized cores as more lateral roots could have found their way into the cores.

## Conclusions

As hypothesized we found that application of the less soil mobile trace elements Cs, Li, Rb and Sr as tracers result in similar uptake rates, whereas the more mobile trace element Se was taken up at a higher rate. As we did not find differences in the background level of the trace elements between species we conclude that the uptake of the less mobile trace elements is affected by RLD in the soil depth where the tracers where placed.

We found that chicory acquired nitrogen from 3.5 m, but did not detect significant uptake of trace element tracers from neither 2.9 nor 3.5 m. However, a substantial increase in tracer uptake in fewer single individual plants indicated that uptake did happen, but our design did not capture this.

We found consistent indications of compensatory deep tracer uptake in chicory intercropped with black medic or ryegrass. Thereto comes that chicory exposed to drought and intercropped with ryegrass resulted in higher tracer uptake per cm of root in the ingrowth cores. Despite that nutrient uptake rate per root length, increases as a response to limited nutrient availability in the topsoil, this is not always enough to counterbalance the lower root length density.

Placing trace element tracers in ingrowth cores did not completely avoid contamination of the soil, but showed to be a feasible design to place trace element tracers as deep as 3.5 m. Modifications of the design such as increasing the number of ingrowth cores would likely improve the ability to use the ingrowth cores in the deeper soil layers.

## Acknowledgments

We thank statisticians Signe Marie Jensen and Helle Sørensen for advice regarding the statistical data analyses and technician Jason Allen Teem for his contribution to the experimental work. All are affiliated with University of Copenhagen. We thank Villum Foundation (DeepFrontier project, grant number VKR023338) for financial support for this study.

